# Unifying genetic association tests via regression: Prospective and retrospective, parametric and non-parametric, and genotype- and allele-based tests

**DOI:** 10.1101/2022.03.31.486648

**Authors:** Lin Zhang, Lei Sun

**Affiliations:** Department of Statistical Sciences, Faculty of Arts and Science, University of Toronto; Division of Biostatistics, Dalla Lana School of Public Health, University of Toronto

**Keywords:** regression, robustness, hypothesis testing, association methods, genome-wide association studies

## Abstract

Genetic association analysis, evaluating the relationship between genetic markers and complex and heritable traits, is the basis of genome-wide association studies. In response, many association tests have been developed, and they are generally classified as prospective vs. retrospective, parametric vs. non-parametric, and genotype- vs. allele-based association tests. While method classification is useful, it is confusing and challenging for practitioners to decide on the ‘optimal’ test to use for their data. Although there are known differences between some of the popular association tests, we provide new results that show the analytical connections between the different tests for both population- and family-based study designs.

**Résumé:** Insérer votre résumé ici. We will supply a French abstract for those authors who can’t prepare it themselves.

## 1. INTRODUCTION

Genetic association analysis, evaluating the relationship between single-nucleotide polymorphisms (SNPs) and complex and heritable traits, is the basis of genome-wide association studies (GWAS). Many association tests thus have been developed, for both population- and family-based study designs (Hayes, 2013; Zhang, 2021).

The development of an association test generally uses one of the following approaches: prospective vs. retrospective, parametric vs. non-parametric, and genotype-based vs. allele-based. While it is useful to classify association tests into different categories, it is confusing and challenging for practitioners to decide on the ‘optimal’ test to use for their data. Differences between some of the popular association tests notwithstanding, the goal of this work is to emphasize the analytical similarities between the different tests, leveraging insights from our recent allele-based association test via regression (Zhang and Sun, 2021).

Before diving into the details of the different association tests, we first briefly review some of the key genetic terminologies necessary for the methodology development and discussion here; for a more extensive list of the statistical genetic terminologies see Table 1 of Zhang (2021), and for biological definitions see Rodriguez et al. (2009).

**Table 1:** Notations for genotype and allele counts and frequency estimates. The HLA-DQ3 marker is an example from Sasieni (1997).

| Genotype Counts and Frequency Estimates |  |  |  | Allele Counts and Frequency Estimates |  |  |
| --- | --- | --- | --- | --- | --- | --- |
| $aa$ | $Aa$ | $AA$ | Total | $a$ | $A$ | Total |
| $n_{aa}$ | $n_{Aa}$ | $n_{AA}$ | $n$ | $n_a = 2n_{aa} + n_{Aa}$ | $n_A = n_{Aa} + 2n_{AA}$ | $2n$ |
| $n_0$ | $n_1$ | $n_2$ | $n$ | $2n_0 + n_1$ | $n_1 + 2n_2$ | $2n$ |
| $\hat{p}_{aa}$ | $\hat{p}_{Aa}$ | $\hat{p}_{AA}$ | 1 | $\hat{p}_a = 1 - \hat{p}_A$ | $\hat{p}_A = n_A/2n$ | 1 |
| $\hat{p}_0 = n_0/n$ | $\hat{p}_1 = n_1/n$ | $\hat{p}_2 = n_2/n$ | 1 | $1 - \hat{p}$ | $\hat{p} = n_A/2n$ | 1 |
| The HLA-DQ3 example from Sasieni (1997) |  |  |  |  |  |  |
| $aa$ | $Aa$ | $AA$ | Total | $a$ | $A$ | Total |
| 313 | 145 | 71 | 529 | 771 | 287 | 1058 |
| 0.59 | 0.27 | 0.13 | 1 | 0.73 | 0.27 | 1 |

We assume the most common scenario where each SNP is bi-allelic with two possible alleles *a* and *A*. By convention, *A* refers to the minor allele with population frequency ≤ 0.5. The corresponding genotype *G* consists of two paired (but unordered) alleles and can be of three possible forms, *aa, Aa* or *AA*.

For a sample of *n* genotypes, we denote the genotypes counts for *aa, Aa* and *AA*, respectively, as *n*_*aa*_, *n*_*Aa*_ and *n*_*AA*_ (or *n*_0_, *n*_1_ and *n*_2_). The corresponding allele counts for *A* and *a* are, respectively, *n*_*A*_ = *n*_*Aa*_ + 2*n*_*AA*_ and *n*_*a*_ = 2*n* − *n*_*A*_. Table 1 summarizes the notations with a data example from Sasieni (1997).

Using *K* alleles instead of 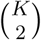 genotypes reduces complexity of a model, but its correct implementation requires the understanding of Hardy-Weinberg equilibrium (HWE). The HWE assumption formulates genotype frequencies as the products of allele frequencies. For a bi-allelic SNP, 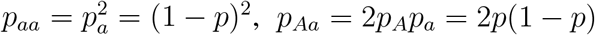 and 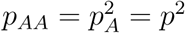, where *p* is also known as the minor allele frequency (MAF). For a SNP to reach HWE, several population-genetics assumptions are needed, including random mating, no inbreeding, infinite population size, discrete generations, and no genetic mutation, migration or selection in a homogeneous population (Lange, 2002).

Departure from HWE or Hardy-Weinberg disequilibrium (HWD) is measured by *δ* = *p*_*AA*_ − *p*^2^ (Weir, 1996). To test for HWD using an independent sample, one typically applies the Pearson goodness-of-fit *χ*^2^ test to the three genotype counts,

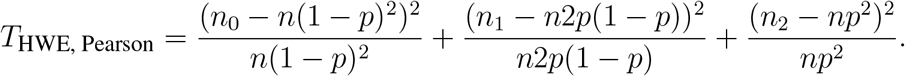

Since *p* is unknown in practice and the allele frequency is replaced by the sample estimate 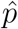, the resulting *T*_HWE, Pearson_ thus follows a 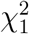 distribution under the null of HWE, reflecting the loss of degrees of freedom (d.f.) from over-fitting the data. Further, *T*_HWE, Pearson_ can be derived from a regression and re-expressed to a form that is directly related to the HWD measure *δ*:

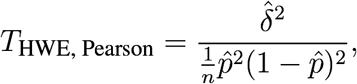

where 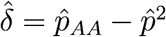 (Zhang and Sun, 2021).

Using the HLA-DQ3 data in Table 1 as an illustration, among a total of 529 individuals, 313, 145 and 71 have genotypes *aa, Aa* and *AA*, respectively. Direct application of *T*_HWE, Pearson_ yields a test statistic of 49.76 and a p-value of 1.74 × 10^*−*12^, suggesting that the SNP is out of HWE or in HWD. As the assumption of HWE may not hold in practice while some association tests explicitly or implicitly assume HWE, our method comparison will pay special attention to the robustness of a method to HWD.

Besides genetic markers, two other main players in genetic association studies are phenotypes (or traits) and covariates (or environmental and clinical factors). As per tradition, we use *Y* to denote a phenotype of interest, which can be binary (e.g. hypertension) or continuous (e.g. height), and we use *Z*’s to denote covariates (e.g. sex and age).

Finally, for a binary outcome, there is the concept of population disease prevalence, *K* = *P* (*Y* = 1). Linking the phenotype with genotypes, there are also the three penetrance probabilities, *f*_0_ = *P* (*Y* = 1|*G* = *aa*), *f*_1_ = *P* (*Y* = 1|*G* = *Aa*) and *f*_2_ = *P* (*Y* = 1|*G* = *AA*).

In the remainder of this paper, we will first review the different classes of tests, namely prospective vs. retrospective, parametric vs. non-parametric, and genotype- vs. allele-based association tests in Section 2. We then present results that show the similarities between the different association tests, 1 d.f. tests in Section 3 and 2 d.f. in Section 4, both for independent samples, which are then followed by tests for pedigree data in Section 5. Finally, we make recommendations for the practical choice of an association test, and we comment on some of the new analytical challenges ahead of us in Section 6.

## 2. A REVIEW OF ASSOCIATION TESTS

Most current genetic association studies adopt what is known as the prospective analytical strategy in the statistical genetics literature, regardless of how the data were collected. In essence, the prospective approach regresses phenotype *Y* on genotype *G* and covariate *Z*. In this context, the genotype *G* of a bi-allelic SNP is typically coded additively as 0, 1 and 2 for the three genotypes *aa, Aa* and *AA* (Hill et al., 2008). The resulting phenotype-genotype association test is the well-known 1 d.f. additive test, based on the parametric phenotype-on-genotype regression.

An alternative to the additive coding is using dummy variables for the three genotypes, which does not impose any assumptions on the genetic effects of the three genotypes on the phenotype of interest. The resulting association test is the well-known but little-used 2 d.f. genotypic test; the 1 d.f. additive model is thought to be the most likely genetic model for most complex traits (Hill et al., 2008), but there are new analytical results supporting the use of a non-additive model (Chen et al., 2021).

The aforementioned two phenotype-genotype association tests, the 1 d.f. additive and 2 d.f. genotypic tests, are both genotype-based tests, regardless of the degrees of freedom. It is not ideal to use the same word genotype to mean two different concepts. But, following the naming convention in the statistical genetics literature, the genotype in ‘phenotype-genotype’ refers to the general data of a genetic marker. The genotype in ‘genotype-based’, on the other hand, refers to how we analyze the genetic data, where the analytical unit is a genotype consisting of two paired alleles (*aa, Aa* or *AA*), in contrast to an allele (*a* or *A*) as in allele-based methods.

That when the three genotypes are coded as 0, 1 and 2 copies of allele *A*, the resulting 1 d.f. additive test is genotype-based is an important but under-appreciated point. Note that the values of 0, 1 and 2, in the prospective phenotype-on-genotype regression, merely specify that the effect of the heterozygous genotype *Aa* on *Y* is the average those of homozygous *aa* and *AA* on *Y* (i.e. additive effect). Like the other genotype-based methods, the 1 d.f. additive test is known to be robust to HWD empirically. We will provide analytical insights on how exactly the prospective, genotype-based, 1 d.f. additive test accounts for HWD in Section 3.

Allele-based (also known as allelic) association tests derived from prospective regressions, however, do not exist to the best our knowledge. The traditional non-parametric allelic test that compares allele frequencies between the case and control groups has been developed (Sasieni, 1997), and it is known to be locally most powerful under certain genetic assumptions including HWE. Subsequently, a robust-to-HWD non-parametric allelic test was developed (Schaid and Jacobsen, 1999). However, before the recent development of an allele-based regression (Zhang and Sun, 2021), the allelic approach is limited to association studies of a binary trait, in an independent sample, and without adjusting for covariate effects. Thus, most published GWAS rely on the prospective phenotype-on-genotype regression and perform a genotype-based association test, typically the 1 d.f. additive test.

In recent years, several authors have considered the so-called retrospective analytical strategy, where the roles of phenotype and genotype are reversed (Thornton and McPeek, 2007; Feng et al., 2011; Feng, 2014; O’Reilly et al., 2012; Jakobsdottir and McPeek, 2013). One motivation for the reverse framework is that most studies sample individuals based on their phenotypic values and then collect their genetic data. Thus, it is statistically preferable to use the retrospective genotype-on-phenotype regression for the purpose of association testing. Another motivation is the need for methods that can analyze multiple (binary and continuous) phenotypes simultaneously for genetic studies of pleiotropy, e.g. the MultiPhen method of O’Reilly et al. (2012). However, the retrospective methods developed so far have not been widely adopted in practice for several reasons, including a lack of clarity about their robustness to HWD.

Back to the popular prospective phenotype-on-genotype regression (with genotypes coded additively), when individuals in a sample are genetically related with each other, using a linear mixed-effect model (LMM) has become the most popular approach for association testing (Eu-Ahsunthornwattana et al., 2014). The variance-covariance matrix of the phenotype is partitioned into a weighted sum of correlation structure due to genetic relatedness and shared environmental effects, where the weight is usually referred to as ‘heritability’ (Visscher et al., 2008). The genetic relatedness is represented by the genetic correlation matrix, from either the known pedigree structure or estimated from the available genome-wide genetic data (Yang et al., 2011; Dimitromanolakis et al., 2019). Although genotype-based association tests are robust to HWD in independent samples (Sasieni, 1997; Schaid and Jacobsen, 1999), little has been discussed in the presence of sample relatedness, which we touch on in Section 5.

The traditional non-parametric allelic test for a binary trait was also extended to study related individuals (Thornton and McPeek, 2007). The test was then generalized to a quasi-likelihood score test to analyze multiple binary or continuous traits (Feng et al., 2011; Feng, 2014). However, none of these methods can directly incorporate covariates. A prospective-retrospective hybrid approach, MASTOR, was then developed to study the association between genotype and one (approximately) normally distributed trait, while adjusting for covariate effects in related individuals (Jakobsdottir and McPeek, 2013). HWE, however, was implicitly assumed by all methods in this category.

Finally, the traditional non-parametric 1 d.f. alellic test can also be derived from the Pearson’s *χ*^2^ test, applied to the 2 × 2 contingency table of allele counts from a case-control study (Table 2). There are also two classic genotype-based non-parametric association tests, the 1 d.f. Cochran-Armitage (CA) trend test and the 2 d.f. Pearson’s *χ*^2^ test, both applied to the 2 × 3 contingency table of genotype counts from a case-control study (Table 2).

**Table 2:** Notations for genotype and allele counts for a case-control study. The HLA-DQ3 example is from Sasieni (1997), studying women with cervical intraepithelial neoplasia 3. The total numbers and notations are the same as in Table 1.

|  | Genotype Counts |  |  |  | Allele Counts |  |  |
| --- | --- | --- | --- | --- | --- | --- | --- |
| | $aa$ | $Aa$ | $AA$ | Total | $a$ | $A$ | Total |
| Case | $r_0$ | $r_1$ | $r_2$ | $r$ | $2r_0 + r_1$ | $r_1 + 2r_2$ | $2r$ |
| Control | $s_0$ | $s_1$ | $s_2$ | $s$ | $2s_0 + s_1$ | $s_1 + 2s_2$ | $2s$ |
| | $n_{aa}$ | $n_{Aa}$ | $n_{AA}$ | $n$ | $n_a$ | $n_A$ | $2n$ |
| Total | $n_0$ | $n_1$ | $n_2$ | $n$ | $2n_0 + n_1$ | $n_1 + 2n_2$ | $2n$ |
| The HLA-DQ3 example from Sasieni (1997) |  |  |  |  |  |  |  |
| Case | 40 | 45 | 28 | 113 | 125 | 101 | 226 |
| Control | 273 | 100 | 43 | 416 | 646 | 186 | 832 |
| Total | 313 | 145 | 71 | 529 | 771 | 287 | 1058 |

In the next few sections, we will show that many of the 1 d.f. (and 2 d.f.) association tests (Table 3) are more similar to than different from each other, particularly when applied to an independent sample of individuals.

**Table 3:**
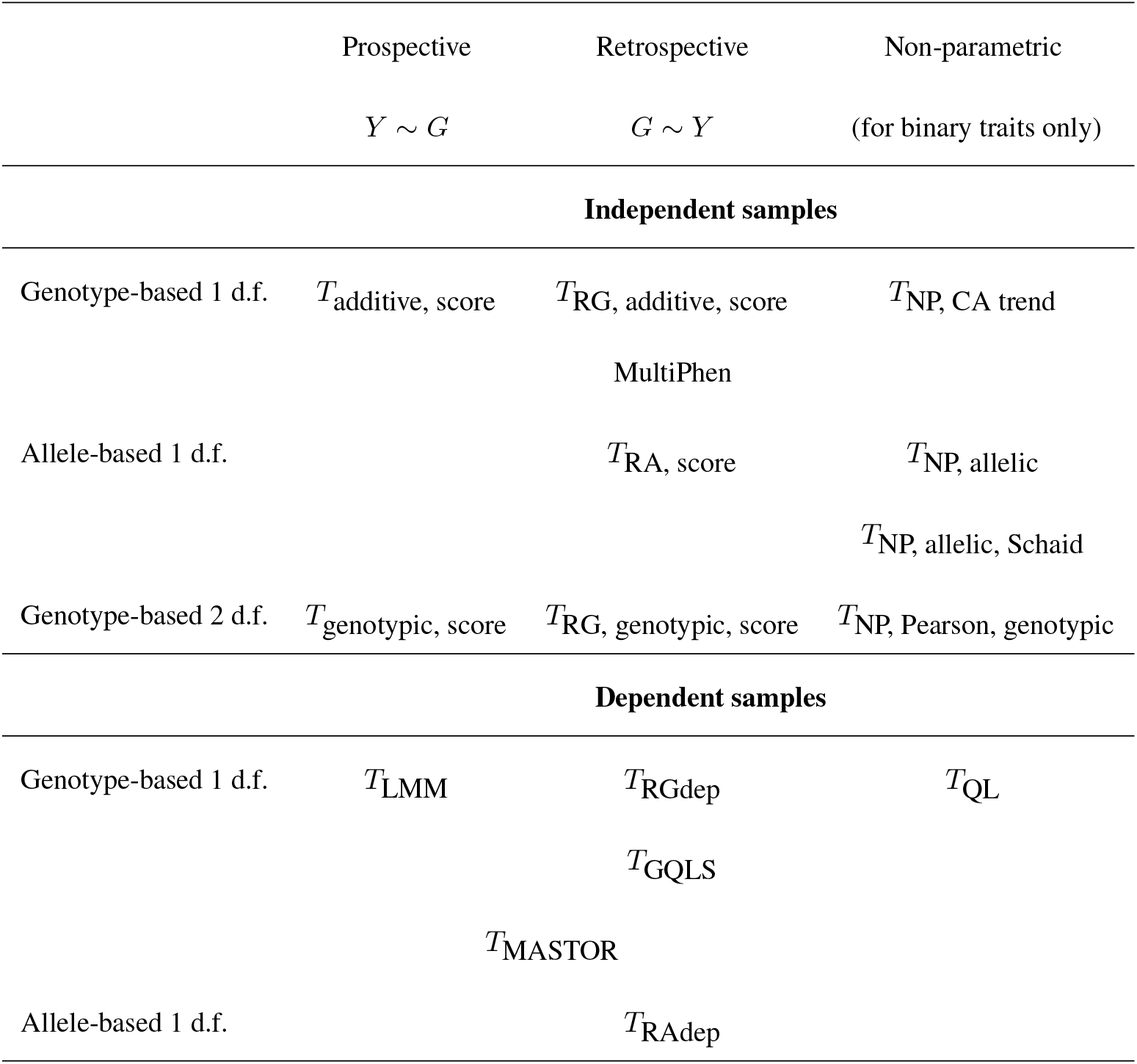
Summary of commonly used genetic association tests. MASTOR is a prospective-retrospective hybrid method. Genotype-based 2 d.f. tests for dependent samples are not discussed here but are possible by using the genotypic coding, instead of the additive coding of *G* = 0, 1 and 2 for the three genotypes, as for independent samples.

## 3. GENETIC ASSOCIATION 1 D.F. TESTS FOR INDEPENDENT SAMPLES

To present the main message with a simple and closed-formed solution, we focus on the simple setting of independent samples without covariates. We show that many of the 1 d.f. score test statistics, derived from prospective vs. retrospective, parametric vs. non-parametric, and genotype- vs. allele-based approaches, have the *exact* analytical form. Along the way, we also propose a new genotype-based retrospective (linear) regression that complements the recent allele-based retrospective regression (Zhang and Sun, 2021). Although the different tests are identical in this simple data scenario, it is easier to use the genotype-based retrospective regression to analyze multiple traits simultaneously.

### 3.1. The traditional prospective 1 d.f. genotype-based additive association test

The most commonly used association test in GWAS, the traditional 1 d.f. additive test is a prospective and genotype-based method that codes the three genotypes *aa, Aa* and *AA* as 0, 1 and 2 respectively, and then uses a generalized linear model (McCullagh and Nelder, 1989). When the phenotype trait *Y* is continuous, the traditional 1 d.f. additive test is derived from the following Gaussian model,

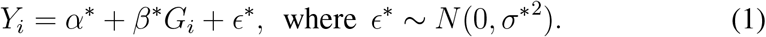

The score test statistic of testing *H*_0_ : *β*\* = 0 is

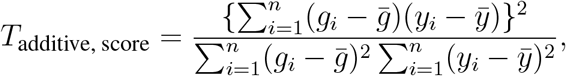

where 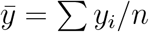 and 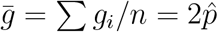.

The derivation of *T*_additive, score_ above is straightforward, but importantly it can be re-expressed as

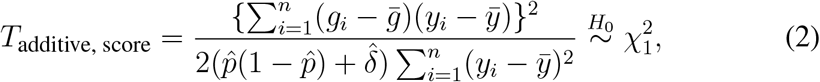

where 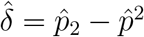 and 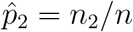. The new expression in (2) helps us understand why the 1 d.f. additive test is robust to HWD: Although the prospective regression (1) does not explicitly model HWD, the resulting test statistic incorporates the HWD estimate, 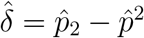, in its variance.

When the phenotype trait is binary, the standard approach is to use the logistic regression,

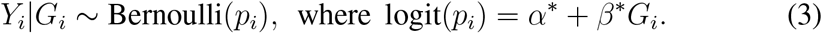

Using the notations in Table 2, the score test statistic of testing *H*_0_ : *β*\* = 0 is

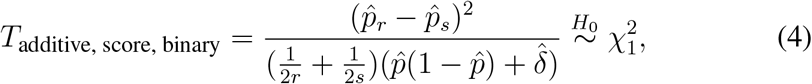

where *r* and *s* are the numbers of cases and controls, respectively. Table 2 shows the additional notations and a data example for a case-control study.

Importantly, when *Y* is binary, *T*_additive, score_ of (2) derived from the linear regression is reduced, *identically*, to *T*_additive, score, binary_ of (4) derived from the logistic regression model. This result is expected as Chen (1983) has shown that, under common regularity conditions, the score test statistics derived from models from the exponential family have *identical* form. In other words, for association testing (not parameter estimation), the Gaussian linear model can be used for a binary outcome. This analytical insight will be utilized later for our proposed retrospective genotype- or allele-based regression models.

### 3.2. Retrospective 1 d.f. genotype- and allele-based association tests

#### 3.2.1. A new retrospective genotype-based (RG) regression

Utilizing the result from Chen (1983), we propose a retrospective genotype-based (RG) regression that simply reverse the role of *Y* and *G* in the above Gaussian model (1). That is, we linearly regress additively coded *G* on *Y*, and co-variates which are omitted for now as noted earlier. This retrospective approach is more flexible for analyzing multiple phenotypes simultaneously, and it will be compared with the Multiphen approach in Section 3.5 and further discussed in Section 5 for analyzing pedigree data of dependent samples.

The proposed RG model is formulated as

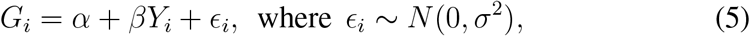

where the *G*_*i*_ is coded additively as in the prospective regression (1). The score test statistic of *H*_0_ : *β* = 0 is then

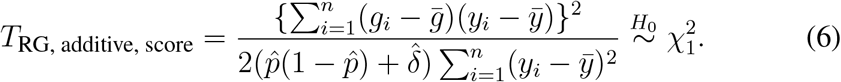

##### Remark 1.

The genotype-based 1 d.f. score test statistics derived from the traditional prospective regression (1) and the proposed retrospective regression (5) are identical: *T*_additive, score_ = *T*_RG, additive, score_. Further, when the trait is binary, the genotype-based 1 d.f. test is testing for allele frequency difference between the case and control groups, but in a robust-to-HWD fashion.

#### 3.2.2. The robust allele-based (RA) regression

We recently developed a robust allele-based (RA) method that is also based on retrospective regression but uses alleles (not additvely coded genotypes) as the response variable (Zhang and Sun, 2021). For an independent sample without covariates, the RA framework is formulated as

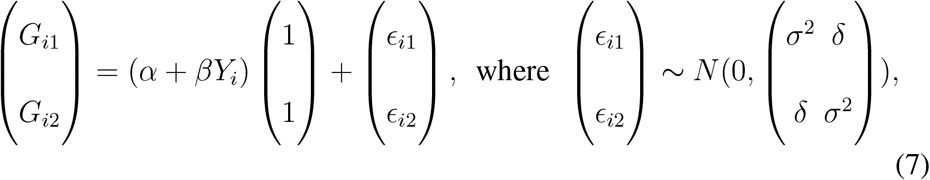

where (*G*_*i*1_, *G*_*i*2_)′ are the two alleles of the genotype of the *i*th individual and

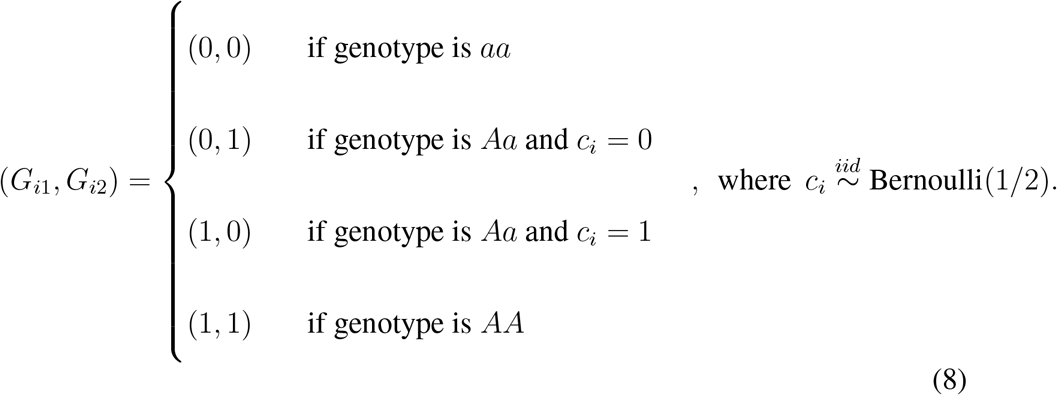

The RA score test statistic of testing *H*_0_ : *β* = 0 is

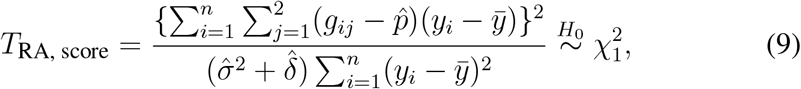

where 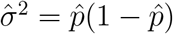 and 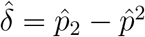. Thus, the RA score test statistic in (9) can be re-expressed as

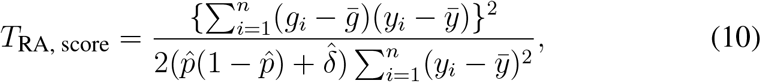

where *g*_*i*_ = *g*_*i*1_ + *g*_*i*2_, which is the number of allele *A* in the *i*th genotype as in (4) and (6).

##### Remark 2.

The 1 d.f. score test derived from the retrospective allele-based regression (7) is identical to the two genotype-based score tests derived from the prospective (1) and retrospective (5) regressions: *T*_additive, score_ = *T*_RG, additive, score_ = *T*_RA, score_. Moreover, this equivalence also holds when adjusting for covariate effects in an independent sample; see Supplementary Material 1 for a detailed proof.

In contrast to the genotype-based retrospective or prospective regressions, the allele-based retrospective regression (7) directly models HWD in the sample using *δ*. Thus, if HWE is assumed (i.e. let *δ* = 0 in the RA model), the resulting score test statistic is

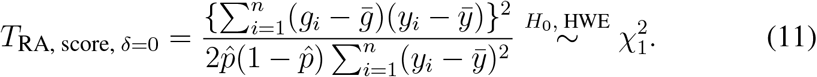

As noted in Zhang and Sun (2021), when *Y* is binary the RA test is further simplified to

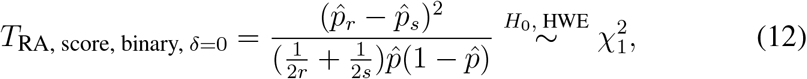

which is the traditional allelic test that is not robust to HWD.

### 3.3. The non-parametric 1 d.f. allele- and genotype-based association tests for case-control studies

#### 3.3.1. Allele-based non-parametric association tests

The traditional allelic test compares the allele frequencies between the case and control groups,

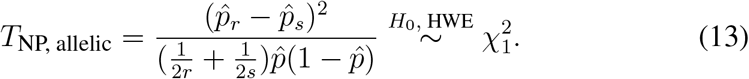

It is easy to show that this test is the same as *T*_NP, Pearson, allelic_, derived from applying the non-parametric Pearson’s 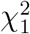 test to the 2 × 2 contingency table of allele counts in Table 2.

The allelic test of (13) is known to be locally most powerful but not robust to HWD. Schaid and Jacobsen (1999) then proposed a robust version by adjusting the variance of the allele frequency difference,

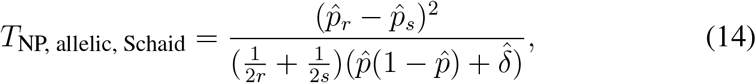

where 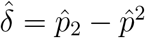 as before.

##### Remark 3.

The traditional non-parametric 1 d.f. allelic test can be derived from the RA regression model, *T*_NP, allelic_ = *T*_NP, Pearson, allelic_ = *T*_RA, score, binary, *δ*=0_, so is the robust version which can also be derived from genotype-based prospective and retrospective regression models, *T*_NP, allelic, Schaid_ = *T*_RA, score, binary_ = *T*_additive, score, binary_ = *T*_RG, additive, score, binary_.

#### 3.3.2. The genotype-based Cochran–Armitage (CA) trend test

The Cochran–Armitage trend test is another non-parametric test for case-control studies. The CA trend test assigns weights *t*_*i*_ = *k, k* = 0, 1, 2, to *aa, Aa* and *AA*, which essentially assumes an additive model for the three genotypes. Thus, it is not surprising to note that the CA trend test has identical form as the other 1 d.f. association tests discussed so far that are also robust to HWD.

Specially, following the notations in Table 2, the CA trend test is constructed based on 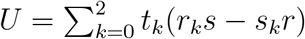 and 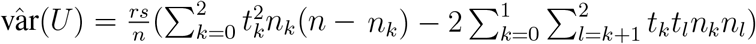. In Supplementary Material 2 we show that

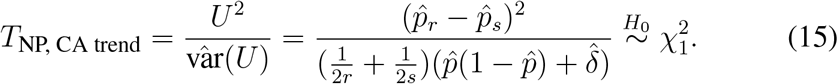

### 3.4. Connections between the 1 d.f. score tests

Figure 1 summarizes the connections between all the aforementioned 1 d.f. association tests that are robust to HWD. We emphasize that all these tests explicitly or implicitly assumes an additive genetic model, and they essentially compare the individual-level phenotype trait value to the pooled sample mean, weighted by the corresponding minor allele counts.

**Figure 1:**
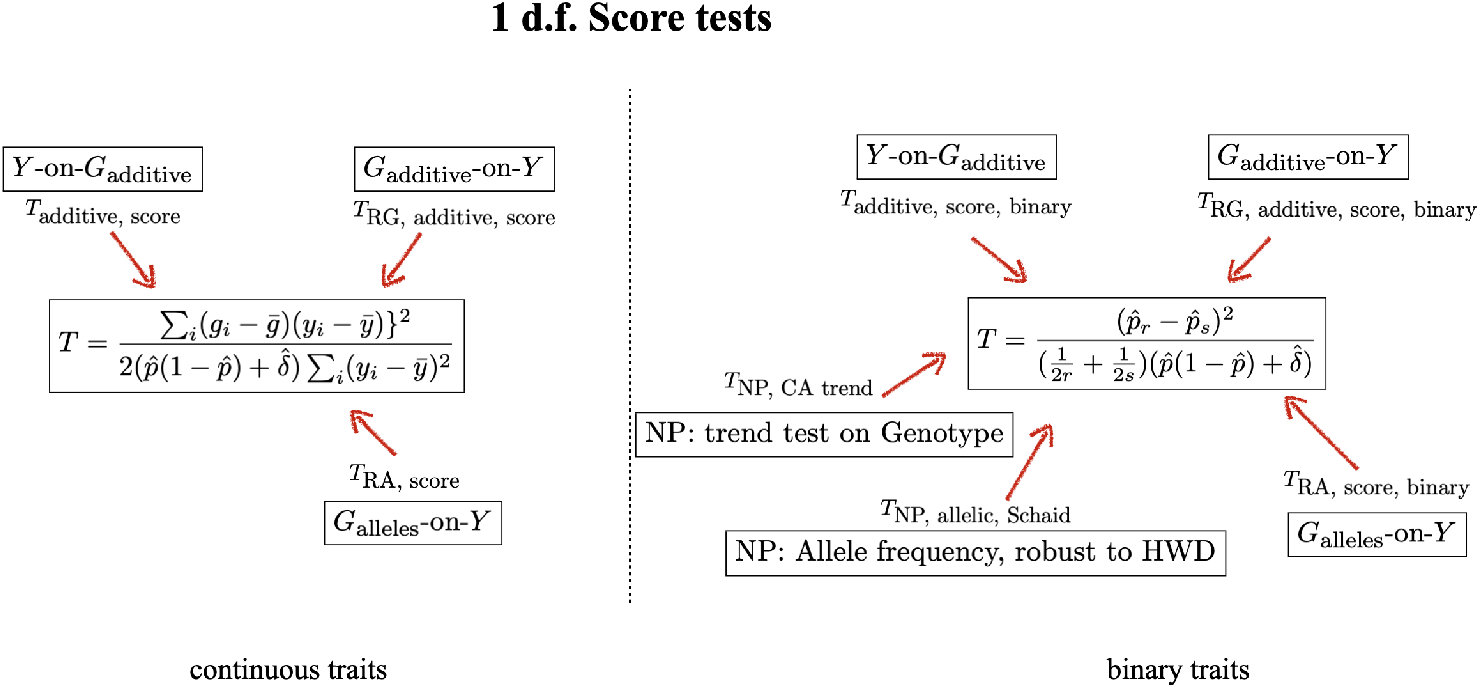
Connections between 1 d.f. additive association tests. *Y* -on-*G*_additive_ refers to the traditional prospective genotype-based regression model that codes the three genotypes, *aa, Aa* and *AA* additively as 0, 1 and 2. NP refers to a non-parametric test. RG refers to the retrospective genotype-based regression in Section 3.2.1, and RA refers to the robust allele-based regression of Zhang and Sun (2021); both RG and RA use *linear* regression. Note that the expression of the test statistic for a continuous trait on the left is analytically reduced to that on the right for a binary trait, which is traditionally derived from the logistic regression.

If the phenotype trait *Y* is binary, another useful insight is that genotype-based tests also compare allele frequencies between the case and control groups but in a robust-to-HWD manner. This robustness is previously known but little understood, but expression in (2) provides the insight on how the most commonly used additive test in GWAS adjusts for HWD through *δ*.

Allele-based tests belong to two groups, robust to HWD or not, but both can be derived by the RA regression model (7). The RA model can also analyze continuous traits unlike the previous allelic tests (Zhang and Sun, 2021). However, if explicit modeling of HWD (e.g. to test for HWD) is not of interest, the retrospective genotype-based regression model (5) is simpler to use as it does not require the partition of the two alleles in each genotype.

### 3.5. MultiPhen

MultiPhen is also a retrospective method that uses genotype-on-phenotype regression but applies an ordinal logistic regression (O’Reilly et al., 2012),

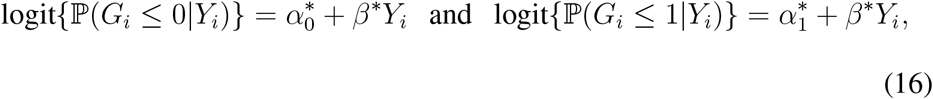

where *G*_*i*_ = 0, 1 and 2 for genotypes *aa, Aa* and *AA*.

By testing *H*_0_ : *β*\* = 0, MultiPhen evaluates the association between *G* and *Y* using a likelihood ratio test (LRT). To obtain a closed form solution and compare it with the other methods, again we focus on the score test,

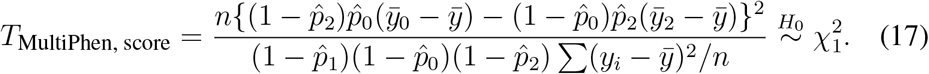

See Supplementary Material 3 for the derivation.

MultiPhen treats each genotype as a category, but it is noticeably as 1 d.f. test, as it assumes a common *β*\* for the ordered genotype categories. To further clarify the analytical nature of Multiphen, we first define

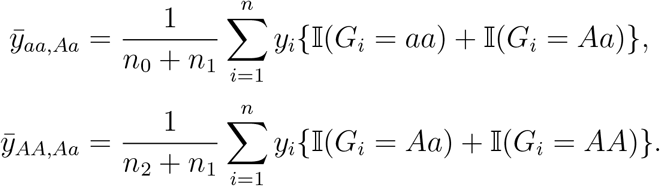

We can then show (Supplementary Material 4) that *T*_MultiPhen, score_ in (17) can be re-expressed as

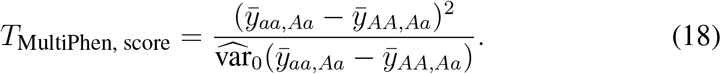

Thus, in essence the 1 d.f. MultiPhen test compares phenotypic means between two combined genotype groups, 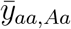 vs. 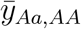. For a binary trait, (18) can be re-expressed as a function of genotype frequencies,

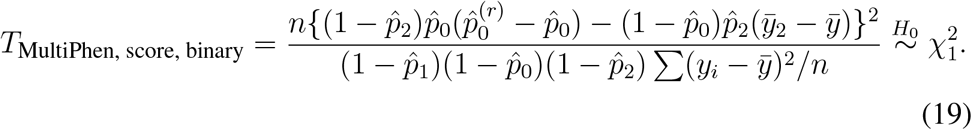

Thus, the Multiphen score test is analytically different from the other 1 d.f. tests shown in Figure 1.

However, similar to the other 1 d.f. tests discussed so far, Multiphen is practically an additive test and may not be able to capture the phenotypic differences in a non-additive model. For example, for a binary disease trait under a heterozygous disadvantage model where *f*_0_ = *f*_2_ = 0.08 and *f*_1_ = 0.1 with risk allele frequency *p* = 0.5, the power of MultiPhen is close to the nominal significance *α* level. This is because, under this non-additive model, the genotype frequencies are (0.2222, 0.5556, 0.2222) for the case population and (0.2527, 0.4945, 0.2527) for the control population, which leads to 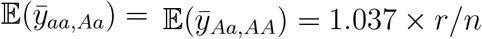, where *r* is the number of cases and *n* is the total sample size.

## 4. GENETIC ASSOCIATION 2 D.F. TESTS FOR INDEPENDENT SAMPLES

The most likely genetic model for complex trait is hypothesized to be additive (Hill et al., 2008). However, as demonstrated above the 1 d.f. additive test may not have power for a heterozygous disadvantage or advantage model, where the penetrance probabilities, *f*_0_, *f*_1_ and *f*_2_, are not monotonically increasing or decreasing across the three genotypes. While the debate on the practical value of a 2 d.f. test is still onging (Chen et al., 2021), we focus on the analytical properties of the various 2 d.f. tests.

Similar to the section above on 1 d.f. tests, we show here that many of the 2 d.f. association tests, derived from genotype-based prospective vs. retrospective and parametric vs. non-parametric approaches have the *exact* analytical form; we discuss a possible allele-based 2 d.f. test in Section 6.

### 4.1. The traditional prospective regression-based 2 d.f. genotypic test

The traditional 2 d.f. genotypic test treats each genotype as a distinct category,

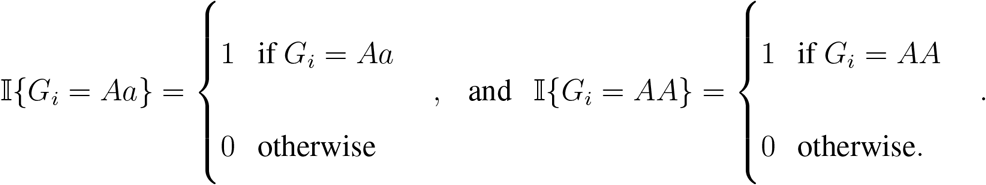

When the phenotype trait *Y* is continuous, the traditional 2 d.f. genotypic test is derived from the following prospective Gaussian model,

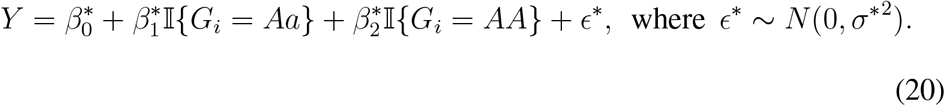

The genotypic test evaluates the association by testing 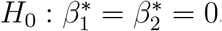, and the corresponding score test statistic is

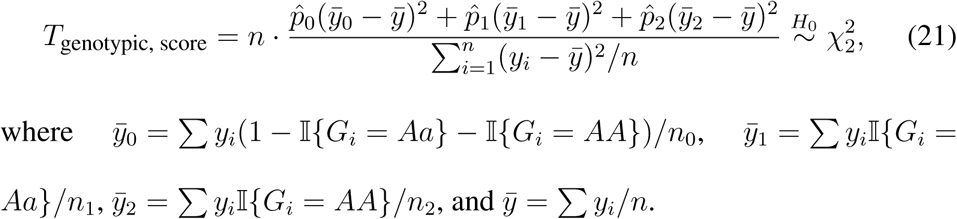

When *Y* is binary, the traditional genotypic model assumes

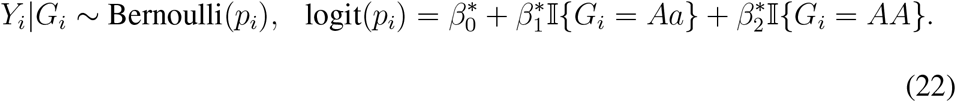

Following the notations in Table 2, we show that the corresponding score test statistic is

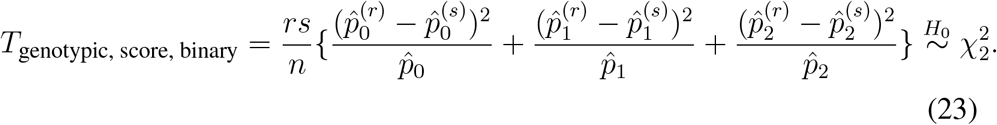

Based on the work of Chen (1983) and the earlier discussion, this test statistic of (23), in its identical form, can also be derived by using a linear model instead of the logistic one in (22). Indeed, when *Y* is binary, *T*_genotypic, score_ in (21) derived from the linear regression can be directly reformulated as *T*_genotypic, score, binary_ in (23) from the logistic regression.

### 4.2. A new retrospective regression-based 2 d.f. genotypic test

The retrospective 1 d.f. genotype-based regression model proposed in Section 3.2.1 codes the three genotypes additively, as in the standard prospective additive model. To arrive at a retrospective 2 d.f. genotypic test, here we first define

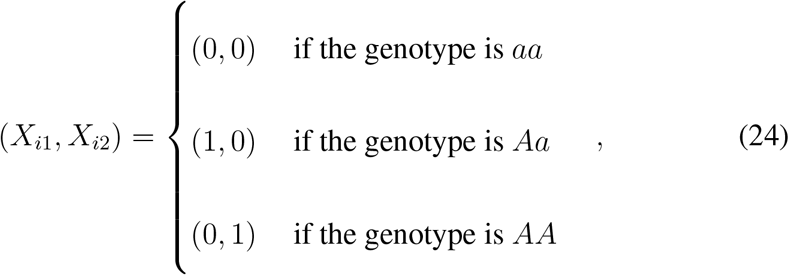

and then specify the retrospective regression model as

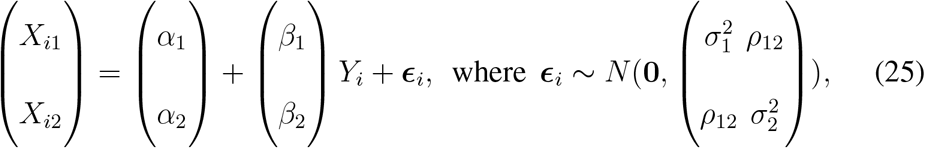

and *ρ*_12_ is the correlation between *X*_*i*1_ and *X*_*i*2_.

Testing association between *G* and *Y* is to evaluate *H*_0_ : *β*_1_ = *β*_2_ = 0, and the corresponding score test statistic is

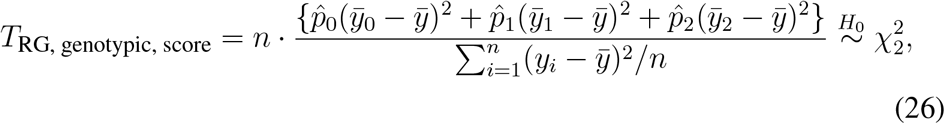

where 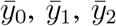 and 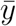 are the same as those defined in (21) for the prospective approach; see Supplementary Material 5. As expressions in (21) and (26) are the same, *T*_RG, genotypic, score, binary_ for a binary trait will be the same with *T*_genotypic, score, binary_ in (23).

### 4.3. The non-parametric 2 d.f. association test for case-control studies

Following the notations in Table 2, it is easy to show that the Pearson’s test is,

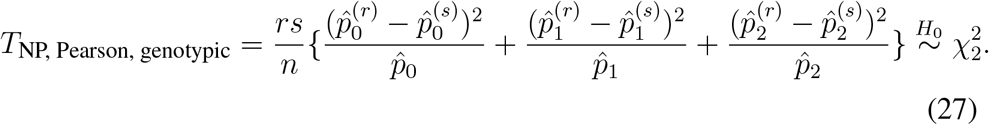

### 4.4. Connections between the 2 d.f. score tests

Figure 2 summarizes the connections between the 2 d.f. tests that treat each genotype as a category. Similar to the 1 d.f. results, the 2 d.f. score test statistics (21) and (26), derived respectively from the standard prospective genotypic model of (20) and the proposed retrospective counterpart (25) have identical form, *T*_genotypic, score_ = *T*_RG, genotypic, score_.

**Figure 2:**
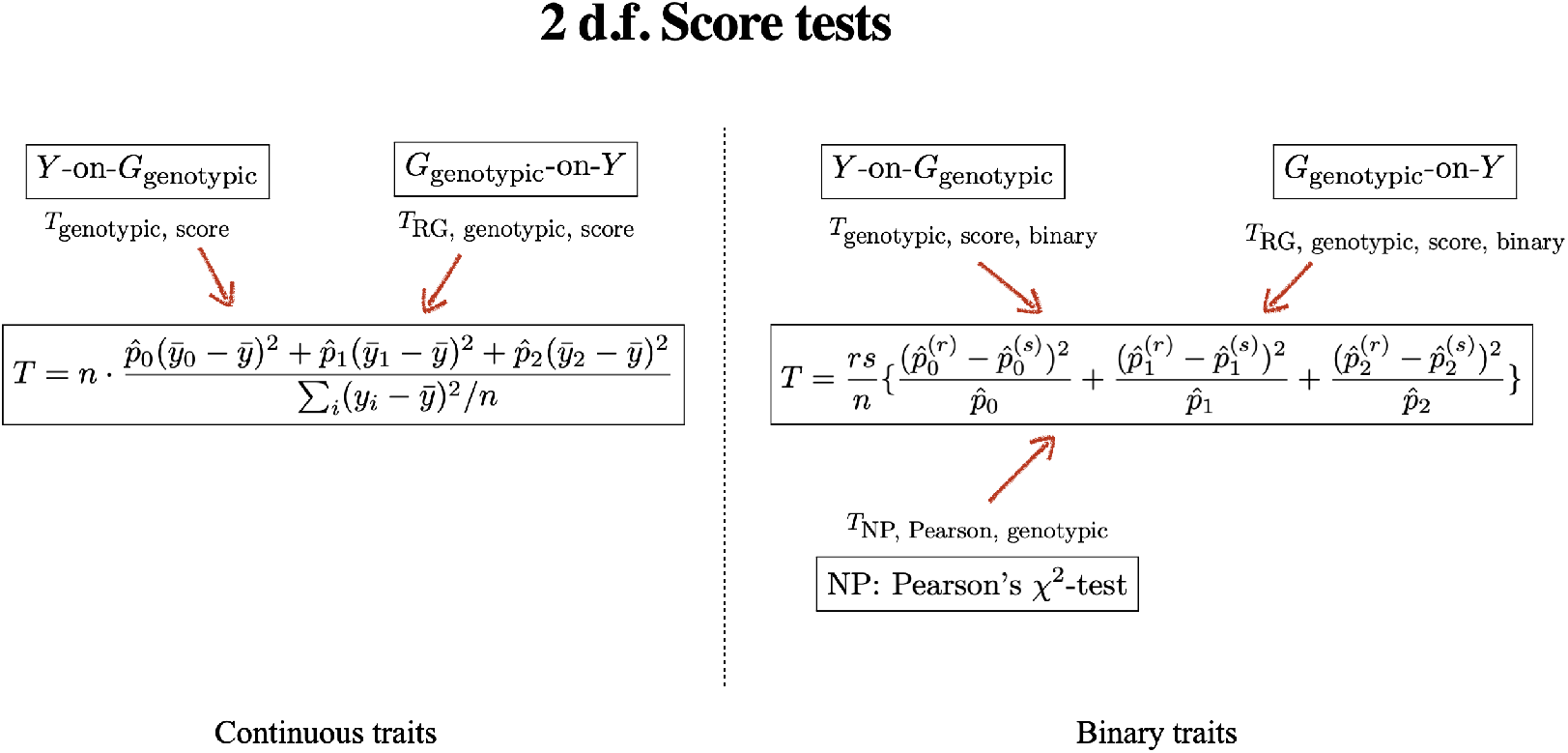
Connections between 2 d.f. additive association tests. *Y* -on-*G*_genotypic_ refers to the traditional 2 d.f. genotypic test that treats genotypes *aa, Aa* and *AA* as categories; NP refers to non-parametric test. RG refers to the retrospective genotype-based regression model in (25), using *linear* regression; see Section 6 for an allele-based 2 d.f. test. Note that the expression of the test statistic for a continuous trait on the left is analytically reduced to that on the right for a binary trait, which is traditionally derived from the logistic regression.

Further, for a binary trait when the non-parametric Pearson’s *χ*^2^ test is applicable, *T*_NP, Pearson, genotypic_ = *T*_genotypic, score, binary_ = *T*_RA joint, ‘score’_ = *T*_RG, genotypic, score, binary_, and they all compare the genotype frequencies between the case and control groups.

## 5. ASSOCIATION TESTS FOR PEDIGREE DATA WITH DEPENDENT SAMPLES

Here we examine the different association tests developed for a sample of genetically related individuals. We still focus on score tests but limit our attention to a) the parametric approach which can model the sample dependence, and b) 1 d.f. tests as the results in the earlier sections suggest that connections made for the different 1 d.f. tests are often generalizable to the 2 d.f. counterparts. We now also explicitly include covariates *Z*’s in a model, and for notation simplicity but without loss of generality, we will use one *Z* to denote multiple covariates unless specified otherwise.

### 5.1. *T*_LMM_, the standard prospective association test from the linear mixed-effect model (LMM)

When individuals in a sample are related to each other as in pedigree data, a common practice is to replace var(***ϵ***\*) = *σ*^*2^*I* in (1) with 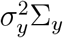 to reflect the sample dependence, and use the linear mixed-effect model,

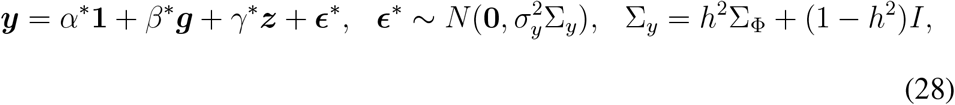

where *h*^2^ is the ‘heritability’, and Σ_Φ_ is the genetic correlation matrix. Among the total variance of 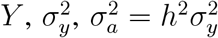 can be interpreted as the variance of *Y* due to additive genetic variation, while 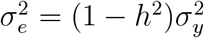 as the variance due to environmental variation.

The score test of testing *H*_0_ : *β*\* = 0 in the LMM of (28) is

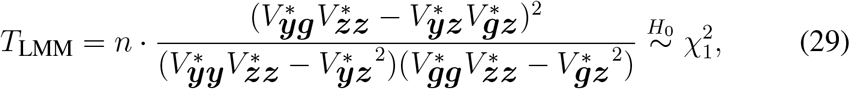

where

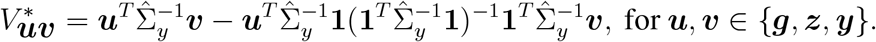

Naturally, when Σ_*y*_ = *I* and *γ*\* = 0 for an independent sample without covariates, *T*_LMM_ in (29) is reduced to *T*_additive, score_ in (2).

For the Σ_Φ_ matrix, Σ_Φ_(*i, j*) = 2*ϕ*_*i,j*_ where *ϕ*_*i,j*_ is the genetic correlation coefficient between individual *i* and individual *j*. However, when the pedigree information is not available, it is a common practice to estimate Σ_Φ_(*i, j*) with the averaged sample correlation across *K* autosomal SNPs (Yang et al., 2011),

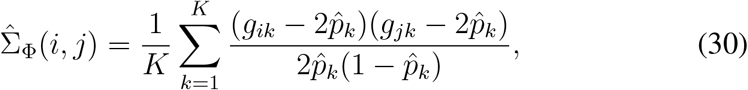

where *g*_*ik*_ and *g*_*jk*_ are the genotypes of SNP *k* for individual *i* and *j* respectively, and 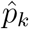 is the estimated allele frequency.

Without the assumption of HWE, var(*G*_*k*_) = 2(*p*_*k*_(1 − *p*_*k*_) + *δ*_*k*_). Thus, (30) implies *δ*_*k*_ = 0, and the traditional genetic correlation coefficient estimation implicitly assumes that HWE holds at each and every SNP *k, k* = 1, …, *K*. Consequently, *T*_LMM_ derived from the linear mixed model (28) is theoretically not robust to HWD, but the practical implication warrants future research.

### 5.2. The retrospective RG and RA regressions for related individuals

In contrast to the standard *Y* -on-*G* prospective approach, where the inference is conditional on *G*, the retrospective alternative can explicit model HWD in *G*. Both the genotype- and allele-based retrospective regression models in Section 3.2 can be extended to adjust for sample dependency.

#### 5.2.1. T_RGdep_ and its connection with T_LMM_

For a sample of related individuals, we extend the retrospective genotype-based regression model in (5) as,

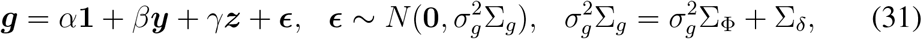

where Σ_Φ_ is the genetic correlation coefficient matrix as defined earlier, and Σ_*δ*_ is a function of *δ*, the HWD parameter. For a pair of siblings for example,

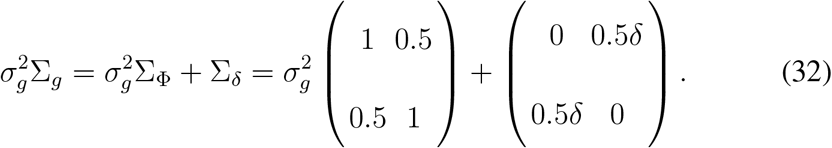

The corresponding score test statistic, testing *H*_0_ : *β* = 0, is

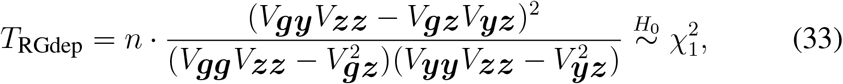

where

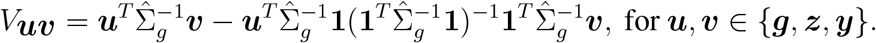

##### Remark 4.

Both expressions in (33) and (29) are symmetric with respect to *G* and *Y*. Thus, if the prospective *T*_LMM_ and retrospective *T*_RGdep_ were to use the same Σ in their respective regression models (28) and (31), then

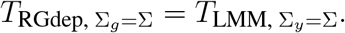

However, the traditional linear mixed-effect model uses genetic correlation coefficient matrix Σ_Φ_ to model Σ_*y*_, while the proposed method adjusts for sample correlation through Σ_*g*_ that includes both Σ_Φ_ for relatedness and a correction factor *δ* in Σ_*δ*_ for potential HWD at the tested SNP.

#### 5.2.2. T_RAdep_ and its connection with T_RGdep_

The retrospective allele-based regression model has been developed for related individuals (Zhang and Lin, 2021), where the variance-covariate matrix in (7) was extended to include the genetic correlation matrix. For the sib-pair case above, it is not surprising to learn that the variance-covariate matrix take the following form,

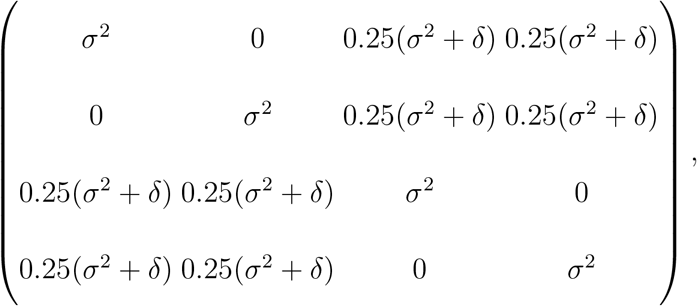

because the RA approach merely ‘expands’ each genotype in the RG model into two alleles. The zero’s in the matrix reflect the fact that the two alleles of an offspring are in HWE after one generation of random mating, even if the parent’s genotypes are out of HWE (Lange, 2002), but HWD can remain between alleles of different siblings (Zhang, 2021).

### 5.3. T_QL_, T_GQLS_ and T_MASTOR_, and their connections with T_RGdep_

#### 5.3.1. T_QL_, the quasi-likelihood-based association test for a binary trait

For a case-control study of a binary trait without covariates and HWD, Thornton and McPeek (2007) proposed an association test that compares sample allele frequency estimates between the case and control groups, while adjusting for relatedness between individuals. The test statistic is defined as,

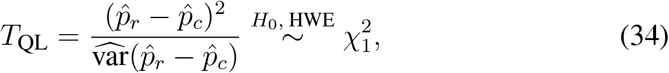

where

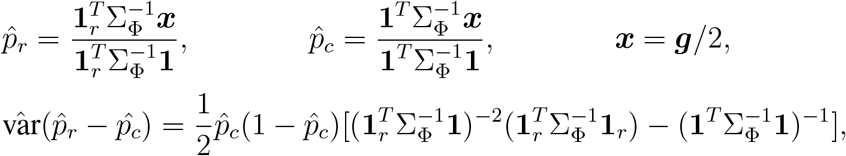

and **1**_*r*_ is a *n* × 1 vector with the *i*th observation to be 1 if individual *i* is from the case group, and 0 otherwise. Note that 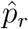 is a sample estimate of the allele frequency in cases, while 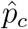 is the pooled estimate using the case-control combined sample, both adjusting for sample correlation through Σ_Φ_.

#### 5.3.2. T_GQLS_, the generalized QL score test

Feng et al. (2011) and Feng (2014) reformulated *T*_QL_ as a generalized quasilikelihood score test that can analyze both continuous and binary traits but still cannot adjust for covariate effects. This quasi-likelihood-based retrospective approach assumes that,

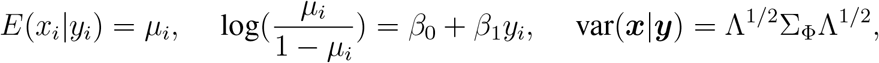

where Λ is an *n* × *n* diagonal matrix, and {diag{Λ}} = {*µ*_1_(1 − *µ*_1_), *µ*_2_(1 − *µ*_2_), …, *µ*_*n*_(1 − *µ*_*n*_)}. Under the null of no association, the framework also assumes that *E*(*x*_*i*_) = *p*_*c*_, and var(***x***) = *p*_*c*_(1 − *p*_*c*_)Σ_Φ_. Thus, the generalized quasi-score statistic of testing *H*_0_ : *β*_1_ = 0 is,

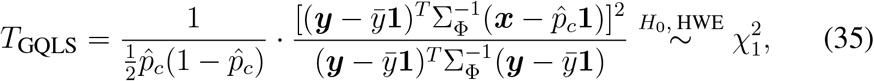

where 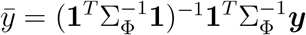, and 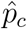 is identical to that in *T*_QL_ of (34). When *Y* is binary, ***y*** = **1**_*r*_. Substituting ***y*** with **1**_*r*_ in (35), it is easy to show that *T*_GQLS, binary_ = *T*_QL_.

##### Remark 5.

When individuals are independent of each other in a sample, Σ_Φ_ = *I*, then 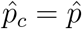. Thus, 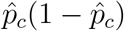 in the denominator of *T*_GQLS_ in (35) assumes HWE when estimating var(*G/*2). In contrast,

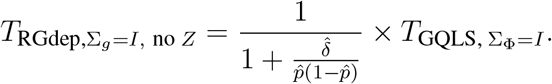

#### 5.3.3. T_MASTOR_, the mixed-effect test for a continuous trait

For a continuous trait with covariates, Jakobsdottir and McPeek (2013) proposed MASTOR, a mixed-effect model-based association test,

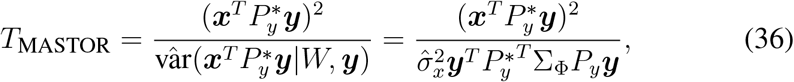

where

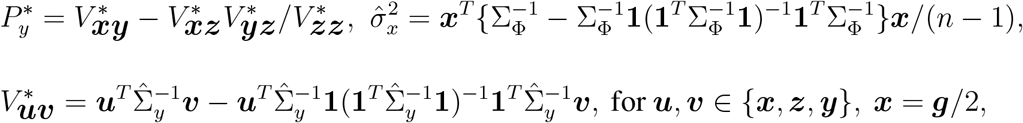

and Σ_*y*_ is the same as that in the linear mixed-effect model of (28), and 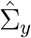 is the maximum likelihood estimate under the null of no association.

To compare *T*_RGdep_ and *T*_MASTOR_, we rewrite *T*_RGdep_ in (33) as

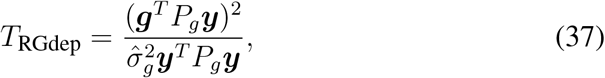

where *P*_*g*_ = *V*_***gy***_ − *V*_***gz***_*V*_***yz***_*/V*_***zz***_, and 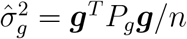. Note that *V*_***uv***_ in (33) (or re-formulated as in (37)) has the same form as 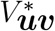 above.

##### Remark 6.

If we were to use Σ_*y*_ = Σ_Φ_ in the calculation of 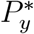; for *T*_MASTOR_ (i.e. assume *h*^2^ = 1 in Σ_*y*_ = *h*^2^Σ_Φ_ + (1 − *h*^2^)*I*), and not accounting for HWD in the calculation of *T*_RGdep_ (i.e. assume *δ* = 0 in 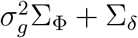), then

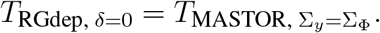

However, although both *T*_MASTOR_ and *T*_RGdep_ measure the correlation between *G* and *Y* while adjusting for covariate effects, the two approaches are distinct from each other. *T*_RGdep_ in (33) is directly derived from a regression framework with *G* as the response variable and *Y* as a covariate, explicitly adjusting for HWD through 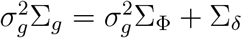 for the tested SNP using model (31). That is, *V*_***uv***_ uses Σ_*g*_ that contains both Σ_Φ_ and Σ_*δ*_ as illustrated in (32) for sibling pairs. In contrast, *T*_MASTOR_ in (36) is a prospective-retrospective hybrid method, where the score function is derived from a LMM with *Y* as the response variable, while the variance of the score function is estimated from modeling *X* = *G/*2 as the response variable. When estimating 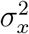 and modelling Σ_*y*_, *T*_MASTOR_ considers Σ_Φ_ alone to account for the genetic relationship between related individuals. Hence, *T*_MASTOR_ implicitly assumes HWE, so do *T*_QL_ and *T*_GQLS_.

## 6. DISCUSSION

We have shown that, for independent samples, many of the score association tests derived from different analytical approaches have *identical* form, as long as the tests have the same degrees of freedom (Figures 1 and 2). The corresponding Wald or LRT tests, derived from the prospective and retrospective models, can differ in finite sample (Zhang 2021), but the equivalency result for score tests here reassures us that the seemingly different prospective, retrospective, genotype-based, allele-based, parametric, and non-parametric tests all lead to the same scientific conclusion.

However, a practical implementation still requires investigators to decide on the d.f. to be used. The 1 d.f. additive test is the most popular test, as the additive genetic model is believed to (approximately) hold for most complex traits. Under an additive model, the 2 d.f. genotypic test loses power as expected. However, recent work has shown that the maximal power loss is capped at 11.4%, regardless of the sample size and nominal significance level (Chen et al., 2021). On the other hand, the 2 d.f. test has good power under a heterozygous advantage or disadvantage genetic model, for which the 1 d.f. may have no power. Given this trade-off, the 2 d.f. test merits consideration in practice. Further, although all the 2 d.f. tests in Section 4 are genotype-based, an allele-based 2 d.f. test is possible for a bi-allelic SNP in a case-control study (Chapter 5 of Zhang (2021)). In that situation, two HWD parameters, *δ*_*r*_ and *δ*_*s*_, can be introduced into the RA model of (7) for cases and controls, respectively. Testing *β* = 0 and *δ*_*r*_ = *δ*_*s*_ is a 2 d.f. test, but a classical score test is ill-defined as *β* = 0 implies *δ*_*r*_ = *δ*_*s*_. The 2 d.f. Wald’s test can be derived, which is not identical to the traditional 2 d.f. genotypic score test, as expected (Chapter 5 of Zhang (2021)).

For dependent samples, the different tests in Section 5 are also connected with each other, *but only if* the same variance-covariance matrix is used. In the standard prospective *Y* -on-*G* regression, the variance-covariance matrix includes the kinship coefficient between the *G*’s even though the inference is conditional on *G*. Further, the kinship coefficient is often estimated from the available data and assumes HWD, which is theoretically incorrect. In contrast, the proposed retrospective *G*-on-*Y* regression includes the HWD parameter *δ* in its modelling of *G*. Another advantage for the retrospective *G*-on-*Y* approach is its ease of analyzing multiple binary and continuous traits.

Adjusting for the main effect of a covariate *Z* can be done in the retrospective regression as shown in Section 5.2.1 and Zhang and Sun (2021), even if its epidemiological interpretation is challenging, as the main purpose of an association analysis is hypothesis testing not parameter estimation. However, including gene-gene (*G* × *G*) or gene-environment (*G* × *Z*) *interaction* into the retrospective framework remains an open problem.

Finally, the work here does not address the challenges associated with analyzing the sex chromosomes, particularly the X chromosome: A male has only one copy of the X chromosome, while a female has two copies. Further, the two copies are subject to random, skewed or no X-inactivation (also known as the dosage compensation), and the true X-inactivation status is unknown. Largely as a result of this model uncertainty, roughly two thirds of GWAS reported by 2013 did not analyze the available X chromosomal data (Wise and Manolio, 2013), and this statistic has been stagnant since then. Recently, just when the problem of X-inactivation uncertainty has finally been solved statistically (Chen et al., 2021), new problems emerged including sex differences in allele frequencies (Wang et al., 2021) and in effect sizes (Bernabeu et al., 2021) between males and females.

In summary, major progresses notwithstanding, developing robust and powerful association methods for *whole-genome* analyses remains an important and exciting topic.

## ACKNOWLEDGEMENTS

This research was funded by the Natural Sciences and Engineering Research Council of Canada (NSERC, RGPIN-04934 and RGPAS-522594). LZ was a trainee and funding recipient of the CANSSI Ontario STAGE (Strategic Training for Advanced Genetic Epidemiology) program at the University of Toronto.

